# Natural Genetic Variation in a Dopamine Receptor is Associated With Variation in Female Fertility in *Drosophila melanogaster*

**DOI:** 10.1101/2022.09.06.506722

**Authors:** Richard F. Lyman, Rachel A. Lyman, Akihiko Yamamoto, Wen Huang, Susan T. Harbison, Shanshan Zhou, Robert R. H. Anholt, Trudy F. C. Mackay

**Affiliations:** Program in Genetics, W. M. Keck Center for Behavioral Biology and Department of Biological Sciences, North Carolina State University, Raleigh NC 27695 USA; Center for Human Genetics and Department of Genetics and Biochemistry, Clemson University, 114 Gregor Mendel Circle, Greenwood, SC 29646 USA; Department of Entomology and Plant Pathology, North Carolina State University, Raleigh NC 27695 USA; Department of Animal Science, Michigan State University, East Lansing, MI 48824; Laboratory of Systems Genetics, Systems Biology Center, National Heart Lung and Blood Institute, Bethesda, MD 20814 USA; Labcorp Drug Development, Morrisville, NC 27560 USA

**Keywords:** diallel cross design, GWAS, RNAi, *Dop2R*

## Abstract

Fertility is a major component of fitness but its genetic architecture remains poorly understood. Using a full diallel cross of 50 *Drosophila melanogaster* Genetic Reference Panel inbred lines with full genome sequences, we found substantial genetic variation in fertility largely attributable to females. We mapped genes associated with variation in female fertility by genome-wide association analysis of common variants in the fly genome. Validation of candidate genes by RNAi knockdown confirmed the role of the dopamine 2-like receptor (*Dop2R*) in promoting egg laying. We replicated the *Dop2R* effect in an independently collected productivity dataset and showed that the effect of the *Dop2R* variant was mediated in part by regulatory gene expression variation. This study demonstrates the strong potential of genome-wide association analysis in this diverse panel of inbred strains and subsequent functional analyses for understanding the genetic architecture of fitness traits.

**Author Summary:** In evolutionary genetics, fitness is defined as the number of offspring an individual contributes to the next generation. This is determined by an individual’s viability (its ability to survive past the reproductive age) and its fertility. Although understanding the genetic basis for natural variation in fitness is a major goal of evolutionary and population genetics, the genetic factors that contribute to variation in fertility in natural populations have remained largely unresolved. To address this issue, we took advantage of the Drosophila Genetic Reference Panel, a population of inbred, sequenced fly lines derived from a natural population. In this panel, there is minimal genetic variation among individuals within each line, whereas variation among the lines reflects the variation observed in the original population from which they were derived. We generated all possible pairwise crosses among 50 of these lines (2,500 distinct genotypes) and measured the productivity (number of offspring produced) for each genotype. We found considerable natural variation in productivity that was primarily determined by the female genotype. We performed a genome wide association as analysis and identified and functionally validated a dopamine receptor that plays a major role in determining variation in female fertility through an effect on egg-laying.

## Introduction

Additive genetic variation for fitness determines the response of populations to natural selection for fitness [1, 2]. The response of a fitness component to natural selection on fitness depends on the additive covariance of the fitness component with fitness, which in turn depends on the heritability of the fitness component and its genetic correlation with fitness [2, 3]. Understanding the magnitude of genetic variation for fitness and its components and the mechanisms maintaining this genetic variation is a major goal of evolutionary quantitative genetics [4, 5]. Individual fitness is defined as the contribution of offspring to the next generation, and is determined by the individual’s viability (survival past reproductive age) and fertility (number of offspring produced). Estimates of genetic variation for viability and fertility in wild populations are confounded by potential genotype by environment correlation and genotype by environment interaction, and the inability to separate male and female effects on fitness. Therefore, much of our empirical knowledge about the genetic basis of variation for fitness and its components is based on laboratory studies, in particular on *Drosophila melanogaster* [6]. The fertility component of fitness is genetically variable in *D. melanogaster* [7-12]; however, the genetic architecture of fertility and the specific genes and variants that contribute to genetic variation in fertility remain poorly understood. Here, we characterize the genetic basis of variation in fertility of young *D. melanogaster* from the sequenced, inbred lines of the *Drosophila* Genetic Reference Panel (DGRP) [13, 14] using a combination of a classical diallel cross design to estimate variance components, high resolution association mapping to identify candidate genes and variants, and RNA interference (RNAi) of candidate genes and variant-based functional validation. Unexpectedly, we find a previously undocumented role of a dopamine receptor affecting natural genetic variation in fertility.

## Results and Discussion

We used a full diallel cross design (reciprocal crosses of parental lines with self crosses included) of 50 DGRP lines to estimate genetic and environmental variance components for fertility. We defined a quantitative trait closely related to fertility of young flies – productivity – as the number of males and females emerging from crosses of four females and four males of the parental lines, with egg laying restricted to 48 hours to minimize larval competition. We measured productivity in three replicate vials for each of the 50 × 50 = 2,500 possible crosses (Figure 1), scoring 737,868 flies in total. We did not find evidence of sex ratio bias (Figure S1), and therefore analyzed the total number of female and male progeny in the 7,500 replicate vials. Approximately 50% of the DGRP lines are infected with the symbiont *Wolbachia pipientis* [14]. We tested the effect of *Wolbachia* infection on productivity and found a small but statistically significant reduction in productivity for crosses when either parent was infected (Table S1, Figure S2). Therefore, we adjusted productivity for the *Wolbachia* infection status of parents before further analyses.

**Figure 1.**
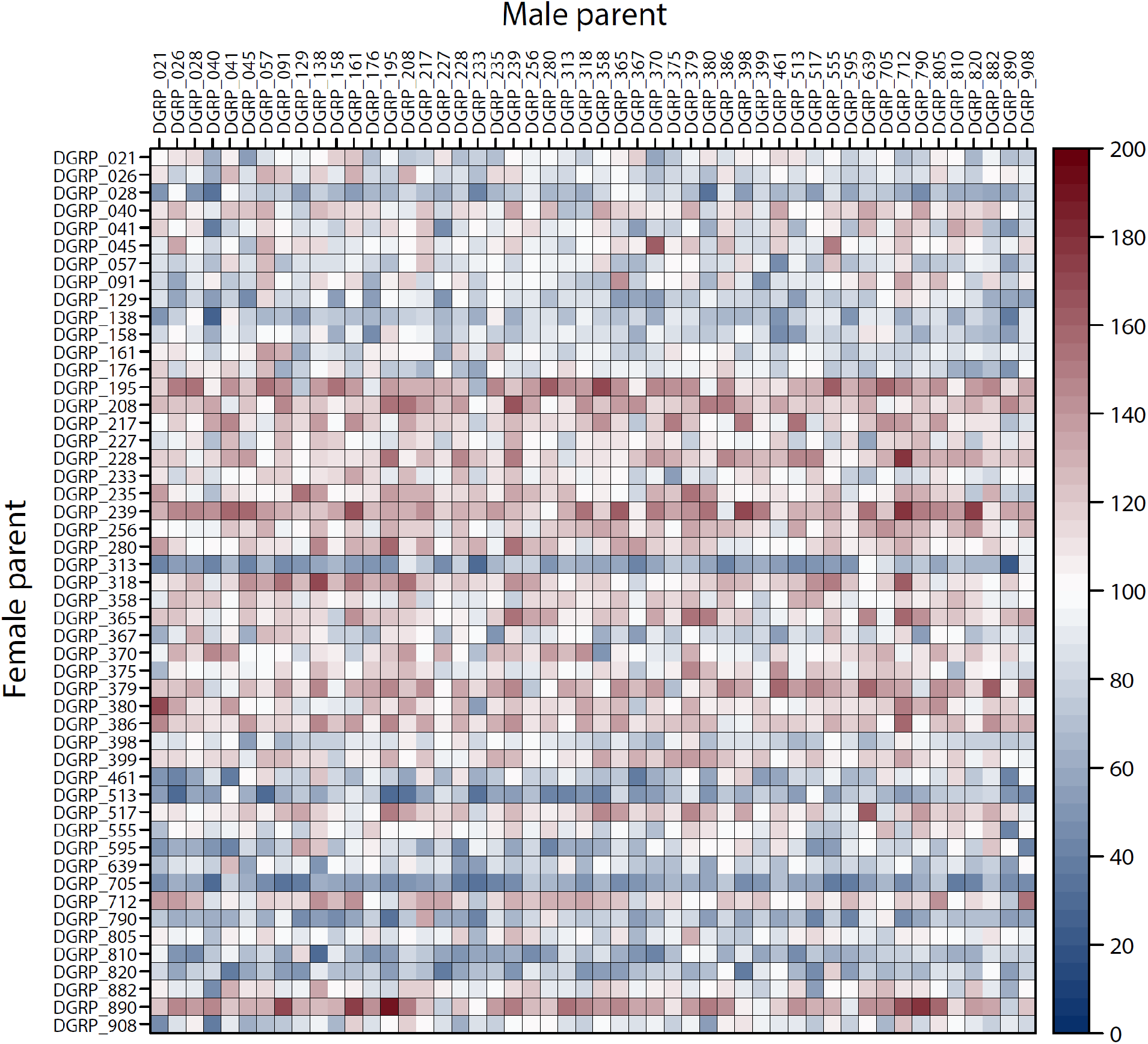
Variation in productivity among crosses of genetically diverse inbred lines. Each cell of the matrix represents one of the 2,500 possible crosses of 50 DGRP inbred lines crossed in a full diallel mating design, with the lines used as sires and dams indicated on the edges of the matrix. The cells are color-coded according to the mean productivity (total number of adult offspring) of three independent replicate crosses, as indicated by the heatmap of productivity.

We observed considerable phenotypic variation in productivity among the crosses. The distribution of productivity appeared symmetrical and normal with a coefficient of variation of 37% (Figure S3). Importantly, 39% of the total variation in productivity was attributable to non-environmental factors, *i*.*e*., parents of the crosses (Table S2). The fact that parents of the crosses differ only by their genotypes implies that the non-environmental variance component is entirely genetic. The genetic variation of productivity observed here includes genetic variation among the female and male parents as well as genetic variation among the progeny. Inbreeding of progeny (from self-crosses) appeared to have negligible effect (*P* = 0.08, Table S3). We further partitioned the genetic variation among crosses according to the bio-model of Cockerham and Weir [15]. This model partitions genetic variance into the additive ‘nuclear’ and ‘extranuclear’ effects of the female and male parents and their two-way non-additive interactions (Figure 2 and Table S4). The nuclear effects are the effects of genes transmitted from parents to offspring that are independent of the sex of the parent. The nuclear effects are only manifested after eggs are fertilized, and are a property of the progeny genotypes. Extranuclear effects refer to any parent-specific effect that is independent of the nuclear genome passed on to the progeny. Extranuclear effects include maternal or paternal genetic effects; parent-of-origin effects of the nuclear genome, such as epigenetic modifications; and maternal cytoplasmic contributions to eggs. Remarkably, our analysis revealed that 85% of the total genetic variation and 33% of the total phenotypic variation in productivity was due to the extranuclear component of the female parents (Figure 2, Table S4). In contrast, variation due to the nuclear genomes of the progeny was not significant.

**Figure 2.**
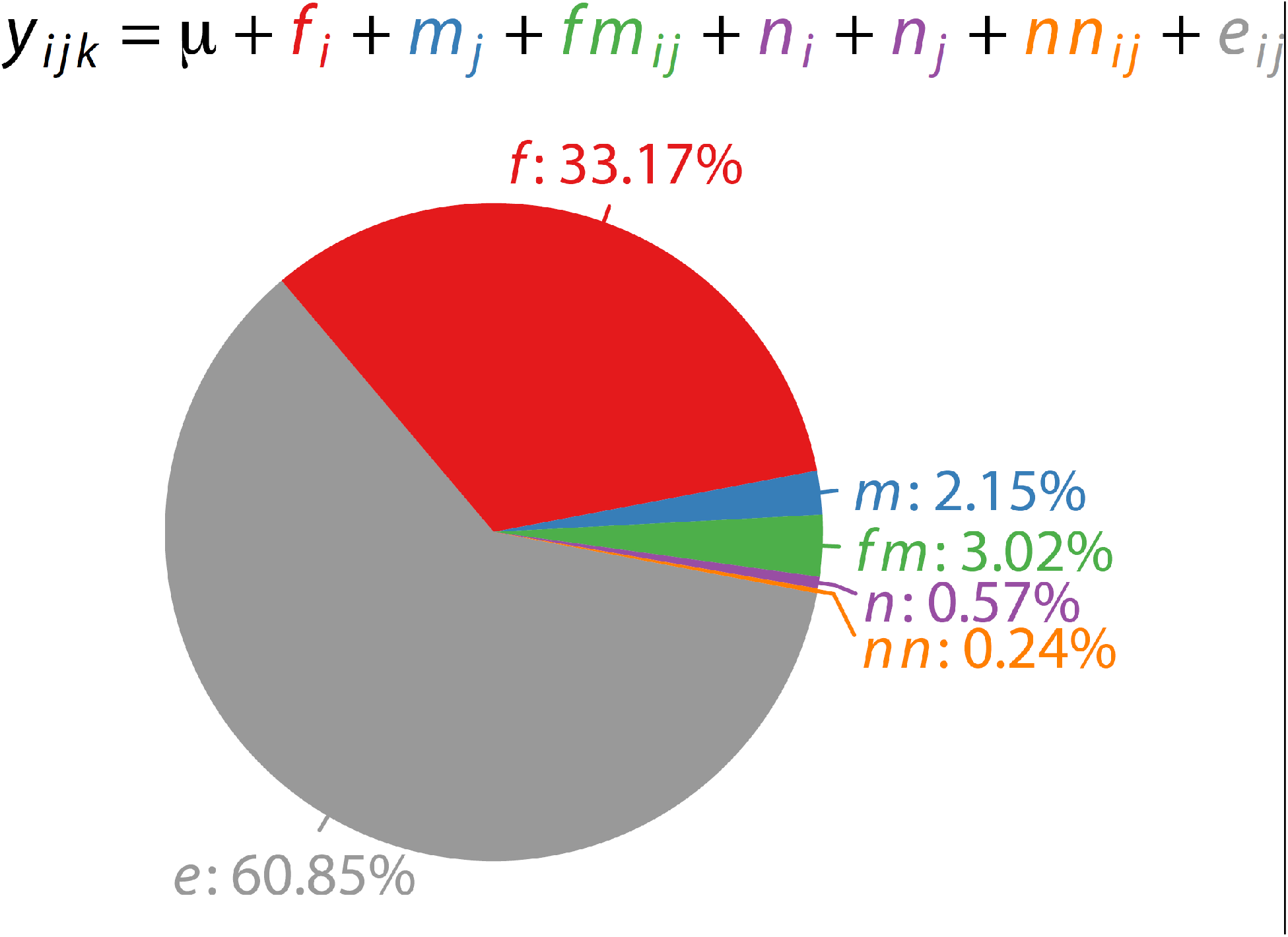
Sources of variation in productivity. The pie chart shows the relative contributions of variance components for productivity from the diallel cross. Variance components were estimated using the mixed effects model *y*_*ijk*_ = *μ* + *f*_*i*_ + *m*_*j*_ + *fm*_*ij*_ + *n*_*i*_ + *n*_*j*_ + *nn*_*ij*_ + *ε*_*ijk*_, where *μ* is the fixed effect for the grand mean; *f, m, fm, n, nn* and *ε* are random effects for the female extranuclear, male extranuclear, extranuclear interaction, nuclear, nuclear interaction, and the residual effects, respectively (see text for explanation); and the subscript for *y* indexes the *i*^th^ female, *j*^th^ male, and *k*^th^ replicate.

We performed a genome-wide association study (GWAS) to identify common (minor allele frequency or MAF ≥ 0.10) single nucleotide polymorphisms (SNPs) associated with variation in productivity of female parents. Among the 1,441,610 SNPs tested, we found a total of 15 with *P*-values < 10^−5^ associated with productivity of females, within or near (within 2kb of the start and end sites) nine genes (Table S5). The most significant SNP (*P* = 3.42 × 10^−7^) is 1,612 bp downstream of the gene encoding the dopamine 2-like receptor (*Dop2R*), and explained approximately 47% of the genetic variance of productivity in females. Female fertility is positively correlated with body size in many insects. Indeed, we found a significant positive correlation between the female component of productivity and female thorax length [16] (*r* = 0.35, *P* = 0.01) but not thorax width (*r* = 0.16, *P* = 0.28). We therefore tested whether the GWAS associations were mediated by the variation in body size by testing for SNP-productivity associations conditional on thorax length. The significance of the tests either remained or diminished very slightly when conditioning the tests on thorax length (Table S5), suggesting that the association cannot be explained by the relationship between fertility and body size.

Given the lenient significance threshold and small sample size (50 DGRP lines), the GWAS is likely prone to false positive associations and overestimation of genetic effects. We therefore obtained additional functional and genetic evidence to corroborate the GWAS results. To validate genes implicated by the significant SNPs in GWAS, we performed two additional analyses. First, we used transgenic *UAS*-RNA interference (RNAi) lines driven by an ubiquitously expressed *GAL4* driver (*actin-GAL4*) to knock down expression of candidate genes in females, and crossed them to males of three DGRP lines with high, moderate, and low male components of productivity. The reduction in scale achieved by focusing on GWAS hits allowed us to further separate productivity into its two components, egg laying and embryonic survival, both of which could potentially be affected by the female parents through extranuclear effects. We therefore counted the number of eggs laid by females and the number of viable adult flies emerging from the crosses. Among the eight genes with publicly available *UAS*-RNAi lines, two (*l(2)37Bb* and *Rpn3*) did not produce viable flies when knocked down. Of the remaining six genes, only knockdown of *Dop2R* in females recapitulated the effects of the SNPs. *Actin-GAL4* > *UAS-Dop2R* RNAi females had significantly reduced productivity compared to the *Actin-GAL4* > RNAi progenitor strain control as well as all other genes (Figure 3a). Furthermore, the effect of *Dop2R* occurred before and/or when eggs were laid because there was also a significant reduction in egg numbers (Figure 3b). In the second analysis, we independently measured the productivity of all (*n* = 204) DGRP females crossed to males of the same lines, including 155 new lines (one line was only present in the diallel experiment). The correlation between the self-crosses in the diallel and this new independent dataset was high (*r* = 0.46, *P* = 8.60 × 10^−4^ for *n* = 49 common lines between the two datasets), testifying to the high heritability and repeatability of the productivity trait. We tested association between the 15 top SNPs from the diallel GWAS and productivity of the self-crosses in the new independent dataset. Remarkably, the *Dop2R* SNP effect remained the most significant one that replicated in this independent dataset of productivity measurement in the DGRP (*P* = 0.0069) (Table S5, Figure 4a).

**Figure 3.**
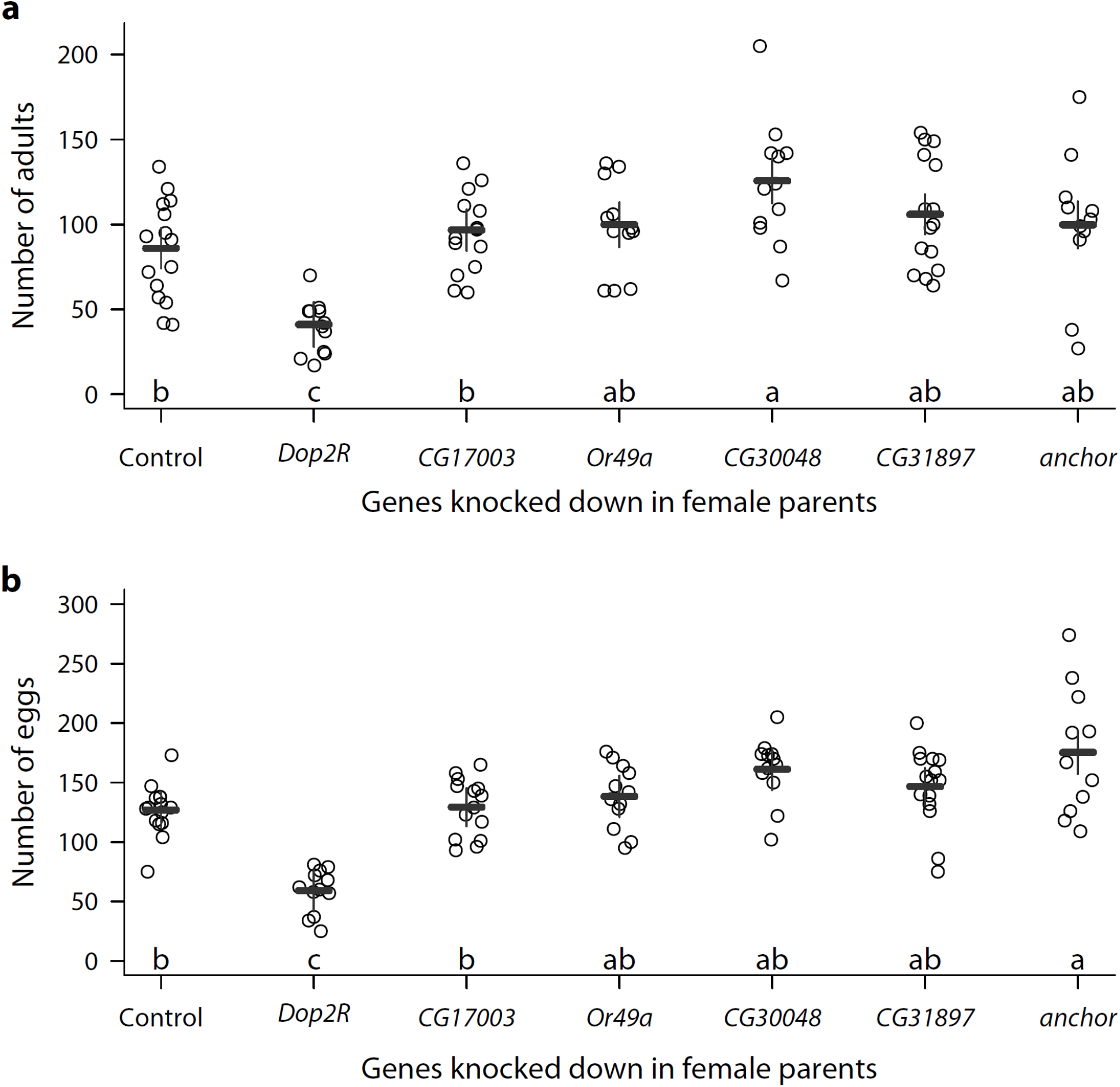
Effects on productivity of RNAi knockdown of candidate genes in female parents. (**a**) The number of adult progeny and (**b**) the number of eggs from *actin-GAL4* > *UAS*-RNAi females for six candidate genes, each crossed to males from representative DGRP lines spanning the range of male genetic variation in productivity. The control is *actin-GAL4* > RNAi progenitor strain females crossed to males of the same three DGRP lines. The horizontal and vertical bars indicate least squares means and standard errors, respectively. Genes with the same letter do not have significantly different numbers of adults or eggs based on Tukey’s multiple comparison at an alpha level of 0.05.

**Figure 4.**
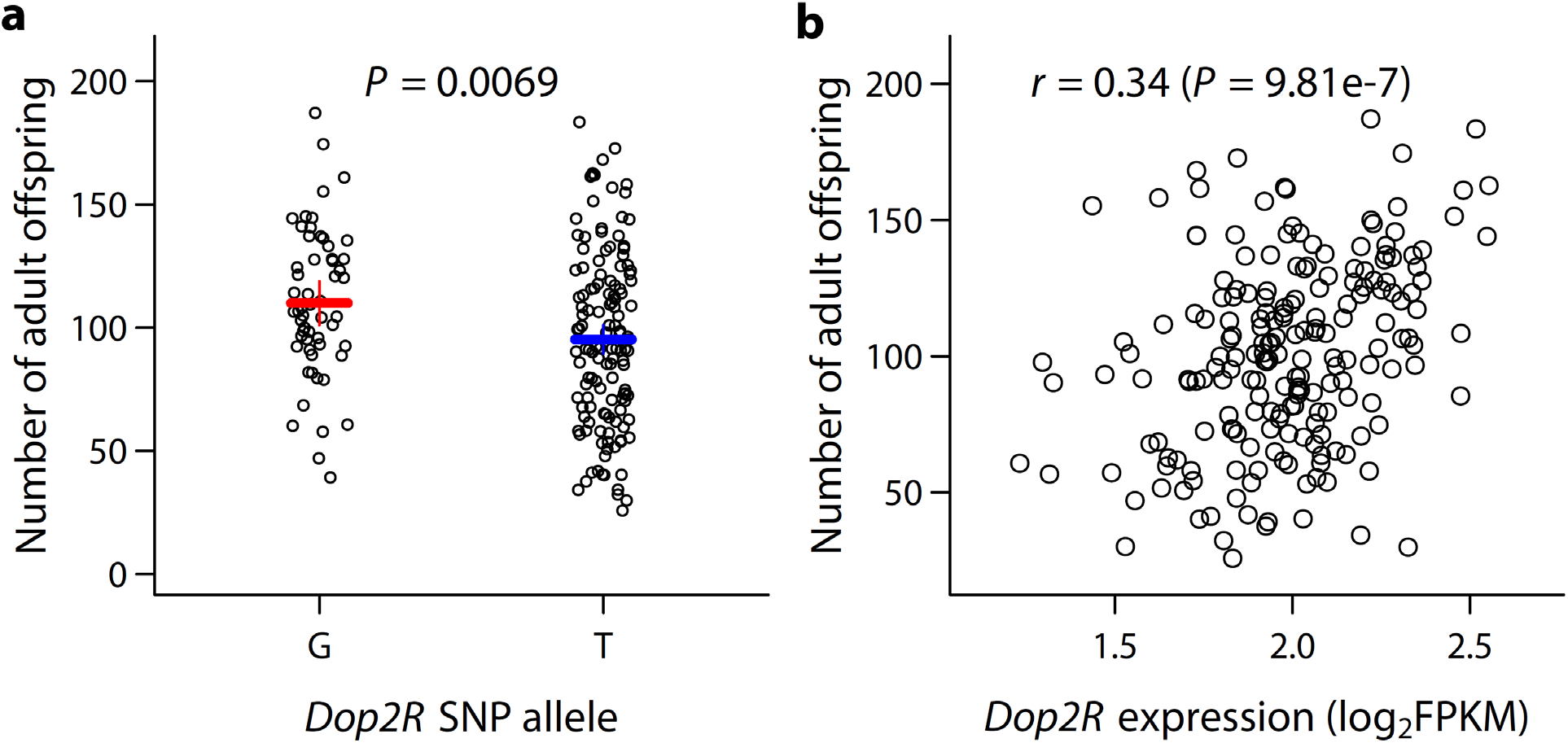
Effects of the *Dop2R* SNP on productivity. (**a**) The number of adult offspring in self-crosses of all DGRP lines according to the *Dop2R* SNP alleles. (**b**) Scatter plot showing the relationship between *Dop2R* expression (log_2_FPKM) and productivity.

*Dop2R* encodes a G-protein-coupled receptor for the biogenic amine dopamine (DA) [17] and is an autoreceptor that regulates the release of dopamine [18]. DA acts as a neurotransmitter, neuromodulator and neurohormone in *D. melanogaster* and other insects. In its role as a neurohormone, DA regulates the levels of juvenile hormone (JH) and 20-hydroxyecdysone (20E), which are required for female oogenesis and fertility; and reduced levels of DA result in abnormal oogenesis and reduced fertility [19-21]. In *D. melanogaster*, pharmacologically decreased levels of DA in sexually mature females (but not males) results in an increase in JH levels [22]. Reduced expression of *Dop2R* in the *corpus allatum* (the endocrine gland in which JH is synthesized) in *D. melanogaster* females using RNAi increases JH levels [23]. Both increases and decreases of JH are associated with defects in oogenesis [22]. Therefore, we hypothesize that the association of *Dop2R* with variation in female productivity in the DGRP may at least partially arise from naturally occurring variation in expression of *Dop2R* in DGRP females, which then causes variation in the amount of DA that regulates fertility via changes in JH and 20E titers. *Dop2R* expression is genetically variable in the DGRP, with a broad sense heritability in females ∼ 0.70 [16]. We found a strong correlation between *Dop2R* expression in females with productivity (*r* = 0.34, *P* = 9.81e-7, Figure 4b) but a relatively weak effect of the *Dop2R* SNP on expression (*P* = 0.09). Mediation analysis revealed that approximately 19% (*P* = 0.09) of the effect of the *Dop2R* SNP on productivity was mediated by the effect of *Dop2R* expression.

It is intriguing that a SNP strongly associated with fertility remained polymorphic in a natural population and the allele increasing fertility was less frequent than the allele decreasing fertility. There are several non-mutually exclusive explanations. First, although there is a very large genetic component of fertility in the laboratory, the genetic variation in the wild may be much smaller compared to environmental variation; therefore the fitness effect is small in nature. Second, this SNP or the causal SNP it tags may be highly pleiotropic, and thus the net fitness effect is small, as would be the case if there is antagonistic pleiotropy with a second fitness trait. Finally, there may be genotype by environment interaction such that the fitness effect varies in fluctuating environments, or at different ages of the flies. This could lead to fluctuation of allele frequency and may allow both alleles to persist.

## Materials and Methods

### Fly husbandry and phenotyping

We arbitrarily selected 50 DGRP lines (DGRP_21, DGRP_26, DGRP_28, DGRP_40, DGRP_41, DGRP_45, DGRP_57, DGRP_91, DGRP_129, DGRP_138, DGRP_158, DGRP_161, DGRP_176, DGRP_195, DGRP_208, DGRP_217, DGRP_227, DGRP_228, DGRP_233, DGRP_235, DGRP_239, DGRP_256, DGRP_280, DGRP_313, DGRP_318, DGRP_358, DGRP_365, DGRP_367, DGRP_370, DGRP_375, DGRP_379, DGRP_380, DGRP_386, DGRP_398, DGRP_399, DGRP_461, DGRP_513, DGRP_517, DGRP_555, DGRP_595, DGRP_639, DGRP_705, DGRP_712, DGRP_790, DGRP_805, DGRP_810, DGRP_820, DGRP_882, DGRP_890, DGRP_908) and performed a full diallel cross, including all reciprocal and within-line crosses. To measure productivity, we placed four virgin females and four males from the respective lines in a vial for two days before removing the adults. We counted the total number of adult females and males up to 16 days post-mating. We performed 3 replicates for each of the 2,500 possible crosses.

### Quantitative genetic analysis

We performed quantitative genetic analyses using SAS PROC MIXED. To test for the effect of *Wolbachia* infection, we fitted a mixed model with *Wolbachia* infection for females and males and their interaction as fixed effects, and line nested within the *Wolbachia* infection as a random effect. We adjusted all observed phenotypic values by the estimated effects of *Wolbachia* infection for subsequent analyses. To partition phenotypic variance, we first fitted a simple model with cross as a random effect. The cross variance component is thus the genetic variance. To partition phenotypic variance according to the bio-model, we computed covariance matrices using relatedness among the crosses such that the covariance between full sibs is 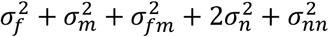, between reciprocal full sibs is 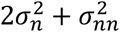, between maternal half sibs is 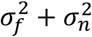, between paternal half sibs is 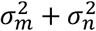, and between reciprocal half sibs is 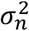; where 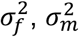, and 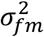 are the variance components for female and male parents’ extranuclear effects and their interaction, respectively; and 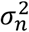 and 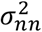 are the variance components for the nuclear genome’s effect and interaction between the two paternal and maternal nuclear genomes, respectively [24].

### SNP calling and GWAS

We used processed alignment files produced by the DGRP Freeze 2 [14] to call SNPs using JGIL [25]. We called SNPs *de novo* to recover sequences for DGRP_398, which was removed from DGRP Freeze 2 because of its relatedness with DGRP_383. We required that SNPs pass a variant level quality score of 500, genotype score of 20, maximum error rate of 0.01 and a minimum allele frequency of 0.1 among the 50 lines. All segregating genotypes within an inbred line were removed from further analysis and thus considered missing. We used the means of productivity across all crosses for the respective female parents as phenotypes for gene mapping. None of the major polymorphic inversions or principal components of the genotypes was significantly associated with the trait. We performed simple linear regressions for GWAS using PLINK [26]. To test for effects of body size, we fitted additionally a model where thorax length was included as a continuous covariate. Gene expression for the *Dop2R* gene was obtained from a previous study in which whole body expression was quantified by RNA-Seq [27]. Mediation analysis was performed using the mediation package in R.

### RNAi and genetic tests for candidate genes

To functionally test candidate genes, we crossed each of the *UAS*-RNAi transgenic lines to the ubiquitous *actin-GAL4* driver line. The *UAS*-RNAi lines were obtained from the Vienna Stock Center, including *CG17003* (*P*{*KK105603*}*VIE-260B*), *Dop2R* (*w*^1118^;*P*{*GD732*}*v11471*), *Or49a* (*w*^1118^;*P*{*GD732*}*v11471*), *CG30048* (*P*{*KK105490*}*VIE-260B*), *DptB* (*P*{*KK112183*}*VIE-260B*), *Rpn3* (*P*{*KK101504*}*VIE-260B*), *l(2)37Bb* (*P*{*KK104117*}*VIE-260B*), *anchor* (*P*{*KK101388*}*VIE-260B*), *CG31897* (*P*{*KK106122*}*VIE-260B*). The adult female offspring (where the respective genes were knocked down) were then crossed to each of three DGRP lines representative of high, medium and low male genetic effects on productivity (DGRP_367, DGRP_375, DGRP_379). The adult flies were transferred 24 hours later to a new vial and visible eggs were counted. The flies in the new vial were cleared another 24 hours later and eggs were counted again. The total number of eggs and adults that emerged were used for subsequent analyses. Five biological replicates for each of the crosses were performed. To independently test the top 15 SNPs from the productivity diallel GWAS, we measured the productivity of self matings for all DGRP lines. We crossed four females and four males of each line in each of 10 replicate vials, and measured productivity as described above for the diallel cross analysis.

### Data and code availability

The raw data and computer codes used in this study are available in a GitHub repository https://github.com/qgg-lab/dgrp-diallel.

## Acknowledgements

We thank the members of the Mackay and Anholt lab groups who generously participated in the Great Fly Count: Gunjan Arya, Tess Brune, Mary Anna Carbone, Kultaran Chohan, Charlene Couch, Kyle Craver, Elizabeth Jones, Katherine Jordan, Faye Lawrence, Lenovia McCoy, Tatiana Morozova, Yazmin Serrano Negron, Elizabeth Ruedi, Lavanya Turlapati, Allison Weber, and Sabu Yamamoto. This work was support by National Institutes of Health grant R01 GM45146 to T.F.C.M and R.R.H.A.

## Supporting Information Captions

**Figure S1. Sex ratios in progeny of the diallel cross**. (**a**) Scatter plot of the numbers of females and males for all 7,500 replicates. The dashed diagonal line indicates equal numbers of females and males. (**b**) Histogram of *P-*values for tests of sex ratio bias. The sex ratio bias for each replicate cross was tested using a *χ*^2^ test for deviation from the expected 0.5 ratio. None of the *P-*values was below 0.05 following adjustment for multiple testing using the Benjamini-Hochberg method.

**Figure S2. Effects of *Wolbachia* infection on productivity**. Boxplots of productivity from the diallel cross, stratified according to infection status of the female and male parents. Statistical tests for significance of the *Wolbachia* effect are given in Table S1.

**Figure S3. Phenotypic variation of productivity**. The histogram depicts the productivity of all replicate crosses. The coefficient of variation (*CV*) of productivity 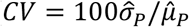, where 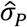 and 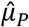 are, respectively, the estimates of the phenotypic standard deviation and the mean productivity) is 37%. The curve is a normal density function with the same mean and variance.

**Table S1. Test for the effect of *Wolbachia* infection on productivity**.

**Table S2. Genetic and environmental variances for productivity among the diallel crosses**.

**Table S3. Test for the effect of inbreeding on productivity**

**Table S4. Sources of genetic and environmental variances for productivity among the diallel crosses**.

**Table S5. Significant GWAS SNPs and implicated genes**.

